# CelFDrive: Artificial Intelligence assisted microscopy for automated detection of rare events

**DOI:** 10.1101/2024.10.17.618897

**Authors:** Scott Brooks, Sara Toral-Pérez, David S. Corcoran, Karl Kilborn, Brian Bodensteiner, Hella Baumann, Nigel J. Burroughs, Andrew D. McAinsh, Till Bretschneider

## Abstract

1

**Summary:** CelFDrive automates high-resolution 3D imaging cells of interest across a variety of fluorescence microscopes, integrating deep learning cell classification from auxiliary low resolution widefield images. CelFDrive enables efficient detection of rare events in large cell populations, such as the onset of cell division, and subsequent rapid switching to 3D imaging modes, increasing the speed for finding cells of interest by an order of magnitude.

**Availability and Implementation:** CelFDrive is available freely for academic purposes at the CelFDrive GitHub repository. and can be installed on Windows, macOS or Linux-based machines with relevant conda environments [1]. To interact with microscopy hardware requires additional software; we use SlideBook software from Intelligent Imaging Innovations (3i), but CelFDrive can be deployed with any microscope control software that can interact with a Python environment. Graphical Processing Units (GPUs) are recommended to increase the speed of application but are not required. On 3i systems the software can be deployed with a range of microscopes including their Lattice LightSheet microscope (LLSM) and spinning disk confocal (SDC).

## 2 Introduction

Imaging living systems with fluorescent microscopy has revolutionised our understanding of a myriad of biological processes by providing information on the dynamics of proteins and cellular structures across multiple time and length scales. The development of ever-improving fluorescent proteins and probes, alongside advances in imaging modalities that can deliver faster, gentler and/or higher resolution imaging continue to advance and deepen our knowledge. One challenge is that cellular events of interest may be rare: for example, a given event may only occur at a specific point-in-time during the cell cycle (e.g. prophase or anaphase) or may only occur when certain conditions are met (e.g. the formation of the immunological synapse when T-cells and anti-gen presenting cells come in close proximity. The reliance on manual identification of such events becomes a major limiting factor in collecting data sets with sufficient power to draw conclusions.

The emergence of automated microscopy methods has the potential to close this gap and accelerate biological discovery. Recent developments within both microscopy and deep learning are enabling imaging systems to “learn” how to identify rare events, then automatically steer the microscope to a given location and collect image sequences. By iterating this loop there is the potential to acquire big data sets of rare events. Approaches have previously been applied to various microscopy methods [2, 3, 4], however, these tend to suffer from phototoxicity and photobleaching effects, which make it difficult to capture dynamic processes at high spatiotemporal resolution. LLSM enables the acquisition of 3D volumes of single cells at high temporal resolution whilst minimising phototoxicity and photobleaching [5]. However, selecting individual cells for imaging is difficult, because a) the field of view is very small (∼ 5-10 *µ*m by ∼100 *µ*m for a cell monolayer) and manual scanning of larger regions of interest is slow and tedious - risking phototoxicty as search times lengthen; b) typical imaging protocols present the microscopist with only a single optical z-section, which furthermore is at an unfamiliar angle with respect to the coverslip (light sheet at 32.8 degrees to the coverslip); c) there is typically only one high magnification objective unlike in many other microscopy methods.

The use of deep learning within bio-image analysis has become increasingly common, from de-noising [6] to segmentation and quantification [7, 8]. This has been facilitated by the emergence of fully convolutional networks [9] which allow various image sizes as input and can benefit from convolutional layers having far fewer trainable parameters than fully connected layers do. In addition, models which can perform classification and localisation at the same time, have made way for real time object detection algorithms [10]. These increasingly advanced architectures and the advent of big datasets have enabled models that can then better generalise to different contexts, enabling us to not necessarily have to retrain models to different contexts [11]. Progress has been made in using deep learning to automate or assist light sheet microscopy capture [12, 13, 14], but many solutions are also difficult to integrate into existing systems and there is not a single software that has been shown to operate effectively across different microscopes, and therefore is not fully utilising the advancements in generalisable deep learning. To address these we have developed CelFDrive, a cross-modality solution to automate LLSM and SDC capture.

## 3 Results & Discussion

### 3.1 Minimal Microscope Modifications for automated LLSM

CelFDrive consists of an automated pipeline to improve the throughput of image acquisition in LLSM. Given the narrow field of view of LLSM, automation requires acquisition of an auxiliary image set to visualise the whole region reachable by LLSM to 1) enable the deep learning, and 2) allow real-time event identification during an experiment.

The commercially available 3i LLSM used here incorporates an air ‘finder’ objective in which the excitation light passes through air before glass at the bottom of the chamber, then through culture media and through the glass coverslip. We performed a low-cost modification to increase the field of view and improve image quality. We replaced the air objective with a Nikon, CFI Plan Fluor 10X, 0.3 NA, water dipping objective with a working distance of 3.5 mm. We replaced the glass bottom with a 0.15 mm thick sheet of fluorinated ethylene propylene (FEP). Between the objective and the FEP we used Immersol W rather than water to reduce evaporation. The FEP is simply held down with grease so that it can be replaced regularly when the chamber is cleaned. Our excitation source was a CoolLEd pE-300-ultra with a far-red excitation filter. The modifications performed improve the speed of acquisition as a 10X objective has a larger field of view, but were not strictly necessary and CelFDrive is compatible with the 3i LLSM imaging setup as it comes.

We held the objective beneath the bath with a custom 3D printed holder made from Polylactic Acid (PLA) that replaced the original 3i objective holder. The increased length of the Nikon 10X objective compared to the original objective required holding the objective barrel/outer-housing rather than using the mounting threads. We secured the objective by tightening a flexure with a M3×24 screw into a brass heat-set insert. The design files are available on the CelFDrive GitHub repository.

### 3.2 Detecting rare mitotic events with Lattice LightSheet Microscopy

We are specifically interested in identifying single cells within a large population that are undergoing (rare) dynamic changes, for example human cells in the prophase stage of mitosis. Prophase constitutes only a few minutes out of a single cell cycle (1440 min) (∼0.3%) and are imminently going to break down their nuclear envelope and initiate key mitotic events i.e. spindle formation, kinetochore assembly, chromosome capture and congression (together termed “prometaphase”). Being able to automate the identification of prophase cells would enable LLSM imaging of nuclear envelope breakdown and the earliest events in prometaphase. However, they are particularly difficult to identify by eye as their nuclear chromatin is somewhat featureless compared to interphase cells.

To enable identification of human RPE1 cells at different mitotic stages, we labeled chromosomes with the membrane permeable dye (SiR-DNA). This emits light in the far-red on excitation in the red, which has benefits in terms of phototoxicity [18]. A large field of view is obtained with a relatively inexpensive inverted 10X magnification 0.3 NA water dipping objective; each image acquired is 550 *µ*m x 850 *µ*m, allowing us to image our whole 2 mm x 2 mm imageable field (Figure 1A). The field of view image is passed to a Python code via SlideBook Conditional Capture and is analyzed by a retrained YOLOv9 [19] neural network to classify cells in six different stages of mitosis (Prophase, Early Prometaphase, Prometaphase, Metaphase, Anaphase and Telophase).

**Figure 1:**
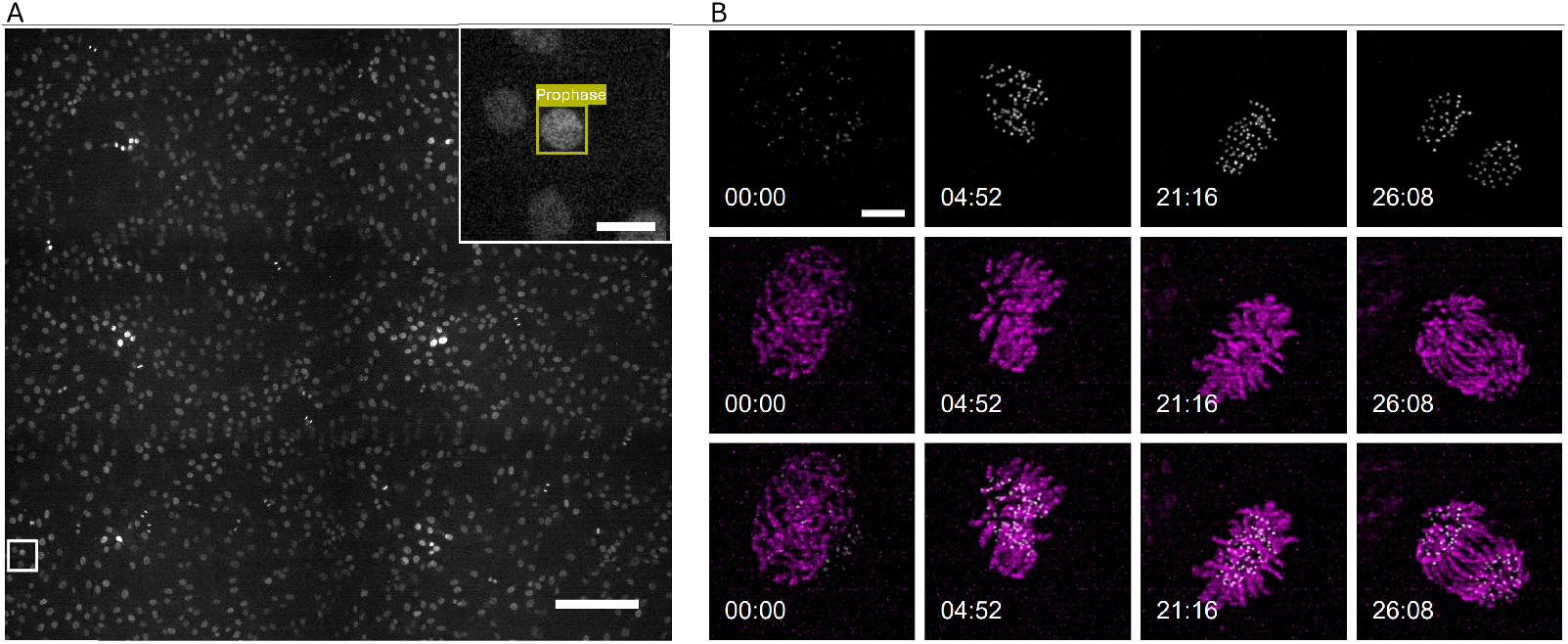
CelFDrive live 3D image acquisition. Human retinal pigment epithelial cells (RPE1) expressing NDC80-EGFP were stained with SiR-DNA to visualise chromosomes and identify cell cycle stages, which are input to the trained YOLOv9 model. A) Montage of widefield fluorescence images of SiR-DNA taken with 10x objective. Montage scale bar length: 200 *µ*m, inset scale bar length: 20 *µ*m. B) The classified prophase cell is then centred on the stage and imaged using LLSM with 20 ms exposure in the 448 nm (white) and 640 nm (magenta) channel for 30 min at 4 s intervals, to capture high spatiotemporal resolution image sequences. Richardson-Lucy [15, 16] deconvolution is performed in skewed space for 10 iterations using a modified version of PetaKit5D[17]. Scale bar length: 5 *µ*m.

The YOLOv9 [19] object detection model was trained using over 18,000 labelled instances of mitotic cells from 4424 images, captured at 10X magnification (Figure 2A).To do this we provide a graphical user interface to rapidly annotate instances of specific cellular states within a time sequence (Figure 2A). CellClicker takes a user drawn bounding box around a region of interest, in our case that of cells in anaphase, where the two sets of separating sister chromatids can be easily identified by their strong fluorescence signal, and opens the image in a new interface. Subsequently, the user moves backwards through the timelapse, simply clicking on the centre of the region of interest, rather than drawing boxes for each instance (Figure 2A). CellSelector then allows to efficiently review cell sequences extracted in this way and for each one associate a label with the earliest timepoint of interest for 3D imaging, such as prometaphase. CellSelector can hold multiple users’ responses which can be aggregated to improve confidence in human annotated data. By providing these interfaces, CelFDrive can easily be retrained and applied to other biological processes.

**Figure 2:**
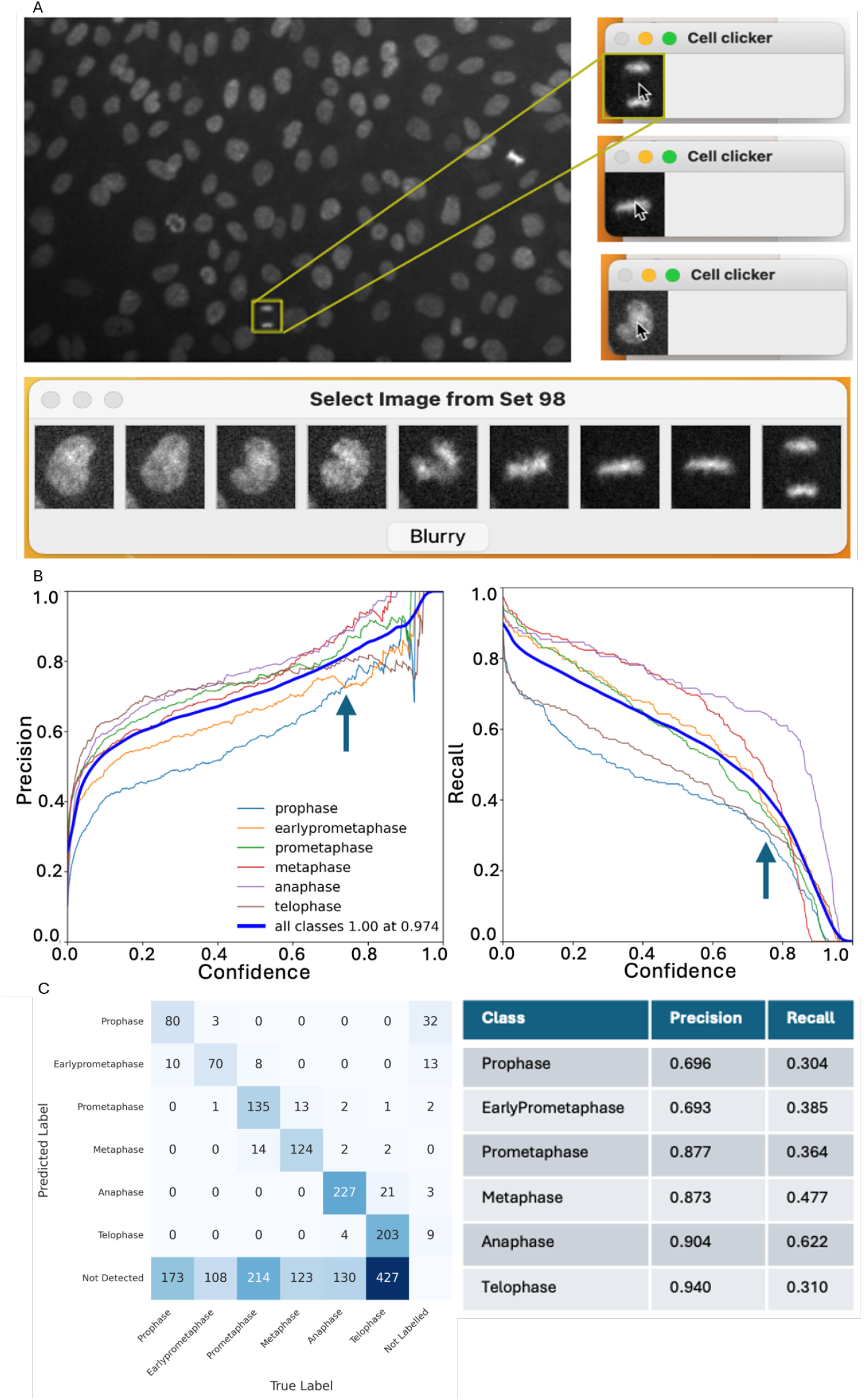
Training network for cell classification A) A bounding box is drawn by the user and the CellClicker UI is opened with the region of interest (ROI). The user clicks on the centre of the cell as it steps back through time until they have reached the start of a biological process. Then the user uses the CellSelector tool to select the phase of interest (e.g. select the timepoint where metaphase occurs). B) Precision-Confidence and Precision-Recall curves for each class, used to determine ideal confidence threshold for the model. A confidence value of 0.75 is selected and shown using an arrow, resulting in lower recall and higher precision. C) Confusion matrix comparing the model predictions versus ground truth on unseen data alongside a table of precision and recall values by class, calculated at 0.75 confidence.

### 3.3 Performance of Object Detection Model

The primary goal was to accurately identify mitotic cells, with a particular focus on prophase. Given the large field of view provided by the low magnification, we prioritised precision over recall, increasing the confidence threshold to 0.75 based on the precision-confidence and recall-confidence curves (Figure 2B). Model confidence in this context is determined by combining box confidence (how likely a predicted box actually contains an object) and class confidence (how confident the model is that the object belongs to that class).

The decision to prioritise precision over recall was informed by the need to reduce false positives and focus on the most confidently detected prophase cells. Opting for increased precision enhances the reliability and throughput of fully automated systems, enabling the generation of large, high-quality datasets. We avoid using accuracy, as it is a poor metric in cases of class imbalance, for example, by classifying all cells as interphase one can achieve an accuracy of approximately 97%.

At 10X magnification, the area on a coverslip accessible for imaging contains approximately 2400 RPE-1 cells, of which around 3% (72 cells) are typically in mitosis, approximately a third of these would be in prophase and early prometaphase. Therefore, despite a reduction in recall (0.304 for prophase), in practice the model still detects a sufficient number of early dividing cells and the model compensates with higher precision (0.696 for prophase, Figure 2C) to ensure when a cell is selected for imaging, there there is a good chance that it is correct in fully automated mode. A user in semi-automated mode is presented with multiple imaging candidates to chose from, allowing them to select for specific characteristics, such as strong expression of a secondary marker.

### 3.4 Imaging rare mitotic events with Lattice Light Sheet Microscopy

The location of the highest scoring cells within the image as classified by the YOLOv9 neural network is converted to stage locations. The stage is then centred automatically on the cell and pre-set LLSM image acquisition parameters are imported. From here, the user can confirm if they wish to begin LLSM live imaging of the cell of interest or select one of the other candidates. An example time-lapse sequence of a cell identified as prophase (Figure 1A, inset) is shown in Figure 1B: In this experiment, RPE1 cells express an endogenous kinetochore marker (NDC80-EGFP, white, top row; [20]) as well as the SiR-DNA, which was used to label chromosomes and identify prophase cells (Magenta, middle row). Kinetochores assemble in a single location on every replicated chromosome (92 per human cell) forming attachment sites for spindle microtubules. These attachments generate the forces to move chromosomes [21]. The initial assembly of the kinetochore (increase in white signal from 00:00 to 04:52) and concurrent condensation of mitotic chromosomes can now be quantified from large numbers of cells immediately after nuclear envelope breakdown (these dynamics and associated biological insights will be reported elsewhere). Furthermore, we can use our automated software [22] to track kinetochore positions over time, providing key information on their dynamic behavior throughout prometaphase. This has proven to be a powerful approach for understanding the behavior of the kinetochore from metaphase to anaphase (e.g. [23] (Figure 1B).

CelFDrive thus opens-up the ability to identify and image rare events just after nuclear envelope breakdown. The software leverages the large field of view provided by the 10X objective, compared to manual scanning where a trained microscopist will take approximately 8 min to find a cell of interest (sample dependent). CelFDrive automation allows a non-technical user to find a cell of interest in under 20 seconds, a twentyfold improvement in speed.

### 3.5 Spinning Disk Confocal

CelFDrive is not limited to LLSM-based applications. We have developed a fully automated prototype using a 3i Marianas spinning disk confocal microscope (Figure 3), which switches automatically between a lower magnification to higher magnification and performs centring to ensure the sample remains within the field of view. For use in fixed cell imaging, we provide a version that montages large regions of the coverslip and images all cells of interest with given criteria. To show the versatility of our model trained on comparably poorer quality images produced by the LLSM’s 10X finder objective, we showed that the network and acquisition algorithm could be applied to an entirely different microscope, changing only the understanding of the coordinates system and introducing a small parfocality (stage direction and pixel spacing). We imaged a montage of 90 locations (1.4 mm x 1.8 mm) (Figure 3A) with a 40X objective (Zeiss, 1.30 NA oil, EC PlnN). After locating mitotic cells of various phases we imaged them with a higher magnification 100X objective to generate high-resolution z-stacks in multiple channels (Zeiss 1.46 NA oil, alphaPlnApo) (Figure 3B). Overall, we were able to screen ∼1000 cells of which 34 were undergoing mitosis. This allows for considerable time gains as it can take about an hour to manually identify and image each coverslip of mitotic cells, and makes it especially powerful in fixed cell immunofluorescence workflows where a user may have multiple coverslips they want to image in a single day. The generalisable nature of CelFDrive also means this can accelerate discovery in almost any cell biology field.

**Figure 3:**
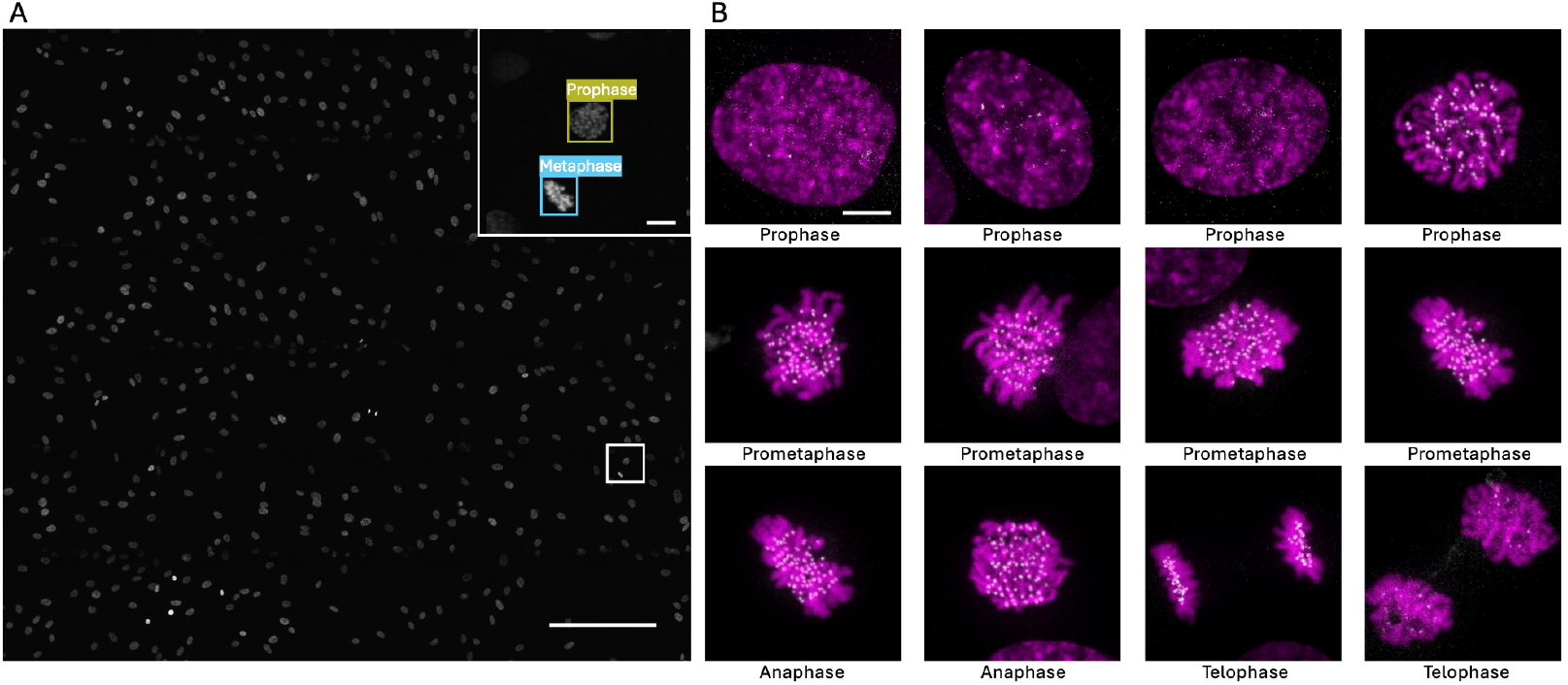
CelFDrive image acquisition of fixed cell samples: A) RPE1 cells expressing NDC80-EGFP were labelled with SiR-DNA to visualise chromosomes and fixed before imaging with spinning disk confocal microscopy. A 2D montage is shown which is stitched together from over 90 images each with a 100 ms exposure in the 640-channel (SiR-DNA) using a 40X objective. Inset shows cell in various cell cycle phases; yellow box, prophase, cyan box, metaphase. Montage scale bar length: 200 *µ*m, inset scale bar length: 10 *µ*m. B) Exemplar two-channel maximum z-projected output images showing various phases of mitosis at 100X magnification, 488 nm (white) and 640 nm (magenta). Scale bar length: 5 *µ*m.

## 4 Conclusions

CelFDrive is a powerful tool that can increase the speed at which expert microscope users are able to find cells in rare states by a factor of 20, increasing the throughput of imaging sessions. CelFDrive is particularly useful for less technical users on the LLSM as it does not require a user to attempt to interpret an optical section of a given cell. We have shown that CelFDrive can be applied to the problem of finding SiR-DNA labelled RPE1 cells in prophase and early prometaphase and provided a model for cell division that should require little retraining from system to system. To assist users in applying CelFDrive to their given biological process, we provide multiple user interfaces to help speed up the process of labelling time-series data. In the future, we will look to fully automate LLSM acquisition and integrate this with essential post-processing such as deskewing and deconvolution. We will also look to expand the training set of the model to take in even more data from other systems, with the aim of creating a generalised model for mitosis classification that does not require retraining from system to system, allowing CelFDrive to become a turnkey application.

## 5 Methods

### 5.1 Cell culture

Human immortilised retinal pigment epitheilal cells (hTERT-RPE1; termed RPE1 throughout) were grown in Dulbecco’s Modified Eagle Medium (F12 medium containing 10% fetal bovine serum (FBS), 2 mM L-glutamine, 100 U/ml penicillin and 100 mg/ml streptomycin) and were maintained at 37ºC with 5% CO_2_ in a humidified incubator. Coverslips were cleaned by immersion in a 1:5 dilution of 37% hydrochloric acid in deionised water and incubated at 65°C for at least 2 h. They were then rinsed twice with deionised water before sonication three times for 5-15 minutes each in deionised water (with three rinses in deionised water between sonication). Coverslips were stored in 70% ethanol with the container sealed with cling film and aluminium foil. Prior to cell plating, coverslips were washed three times with PBS, ensuring complete removal of residual PBS by aspiration. For Lattice LightSheet microscopy, cells were seeded at 90% confluency on pre-cleaned 5 mm coverslips within a 35 mm tissue culture plate. For confocal microscopy, cells were seeded at 90% confluency on pre-cleaned 22×22 mm coverslips placed in 6-well plates.

### 5.2 Fixation

Following a 1-hour incubation with 0.5 *µ*M SiR-DNA [18], the cells were fixed using PTEMF buffer (20 mM PIPES pH 6.8, 0.2% Triton X-100, 10 mM EGTA pH 7.0, 1 mM MgCl_2_, 4% formaldehyde in double-distilled H_2_O). To fix the cells, the culture medium was aspirated, and 2ml of PTEMF fixative was added to each well, followed by a 10-minute incubation at room temperature. After fixation, the fixative was removed, and the cells were washed three times with PBS with gentle rocking for 10 min. Finally, the coverslips were mounted onto slides using vectashield (Vector Laboratories, Inc., Burlingame, CA, USA) and sealed with nail varnish.

### 5.3 Imaging

Imaging of live cells was carried out using a Lattice LightSheet microscope [5] which was manufactured by 3i (https://www.intelligent-imaging.com), with a Coherent Sapphire 300mW 488nm laser and a MPB Communications Inc 500mW 642 laser. The objectives were a Special Optics 0.7 NA LWD WI for excitation and a Nikon CFI Apo LWD 25x 1.1 NA WI for detection. The two cameras were Hamamatsu ORCA-Flash4.0 V3. The acquisition was controlled with SlideBook 2024.2.

Following a 1-hour incubation with 0.5 *µ*M SiR-DNA[18], the coverslip was mounted onto a sample holder using grease and then transferred to the LLSM bath containing CO_2_-independent Leibovitz’s L-15 medium. Prior to each imaging session, the LLSM light path was aligned. Subsequently, bead images were acquired for each imaging channel (488 nm and 640 nm) and used for deconvolution.

Images from the ‘finder’ objective were captured using an inverted 10X Nikon, CFI Plan Fluor 10X, 0.3 NA, water dipping objective with a Hamamatsu ORCA-spark Digital CMOS camera. The excitation source was a CoolLEd pE-300ultra, which was filtered for far red excitation. For each coverslip imaged, the finder objective was aligned with the LLSM objectives to ensure that the cell centered in focus with the finder objective corresponded to the cell currently in view with the LLSM detection objective. A 2×3 montage covering an area of 1.7mm x 1.65mm within the 2mm x 2mm field accessible by the LLSM was captured at 18% LED neutral density (ND). The montage was then passed to CelFDrive to detect mitotic cells, before centering the selected cell of interest for LLSM imaging.

Multichannel 3D time-lapse images of Ndc80-eGFP were acquired with 8% laser power (30 mW) and the 640 nm channel with 1% laser power (50 mW). Volumes were collected in sample scan mode with an exposure time of 20 ms per slice with 60 planes and a 0.5 *µ*m Z-step (corresponding to a 0.271 *µ*m Z-spacing after deskewing). Each stack took 2.7 s to acquire, and there was a 4 s interval between the start of each timepoint, allowing a 1.3 s rest period between timepoints to minimise phototoxicity. The acquired time-lapses were deconvolved [15] [16] in skewed space using 10 iterations of the Richardson-Lucy algorithm, then deskewed, with both processes performed using a modified version of PetaKit5D [17] (https://github.com/warwickcamdu/PetaKit5DfromSLD), which automates the workflow from the original SlideBook files to the final output in TIFF format.

Imaging of fixed cells was carried out using a 3i Marianas spinning disk confocal equipped with 2× Photometrics Prime 95B sCMOS cameras. The initial montage was acquired using a 40X oil immersion objective (1.30 NA, EC Plan-Neofluar, Zeiss) to scan a large area, while high-resolution imaging was conducted with a 100X oil immersion objective (1.46 NA, alpha Plan-Apochromat, Zeiss). To account for the coverslip tilt, the initial montage was defined by focusing on 3 extent points. Parfocality adjustments were calculated by subtracting the difference between the Z focal positions when switching between the high and low magnification objectives, with this adjustment provided to CelFDrive. The low magnification montage was captured in 3D using the 640 nm channel with 2% laser power at 50 ms exposure time per optical section, with a Z-spacing of 2 *µ*m across 21 planes, to account for the different Z positions of mitotic cells. The 3D montage was then maximum intensity projected (MIP) in SlideBook’s scripting language before being transferred to CelFDrive for object detection and coordinate calculation. The output locations were imaged with the 40X objective using a 488 nm laser at 100% power and 640 nm laser at 1% power. The exposure time for both channels was set at 100 ms per optical section, with a Z-spacing of 0.5 *µ*m across 41 planes.

### 5.4 Software

CelFDrive can be installed by downloading the GitHub repository and installing the conda environment provided [1]. The primary code is written in Python [24] and the model is retrained using the Ultralytics Python package [25] from a pretrained YoloV9 [19] model for 100 epochs. The software is free to use for academic purposes, but will need to integrate with a microscope control software that has the ability to interact with Python. We used 3i’s SlideBook software’s conditional capture and scripting language.

## 6 Author contributions

Scott Brooks: Conceptualisation, Data Curation, Formal Analysis, Investigation, Methodology, Software, Visualisation, Writing – Original Draft Sara Toral-Pérez: Data Curation, Methodology, Investigation, Validation, Visualisation, Writing – Original Draft David S. Corcoran: Data Curation, Methodology, Investigation, Writing – Original Draft Karl Kilborn: Methodology, Software, Writing – Reviewing & Editing Brian Bodensteiner: Methodology, Software Hella Baumann: Supervision, Methodology, Writing – Reviewing & Editing Nigel J. Burroughs: Supervision, Conceptualisation, Writing – Reviewing & Editing Andrew D. McAinsh: Supervision, Conceptualisation, Funding Acquisition, Methodology, Resources, Writing – Original Draft, Writing – Reviewing & Editing Till Bretschneider: Supervision, Conceptualisation, Funding Acquisition, Methodology, Resources, Writing – Original Draft, Writing – Reviewing & Editing

## 7 Acknowledgements

We would like to thank Warwick’s Computational and Advanced Microscopy Development Unit (CAMDU) for their continued support, providing training, maintaining microscopes and analysis machines. At our industry partner, 3i, we would like to thank Owen Richards for his work with technical issues relating to LLSM and Benjamin Atkinson for helping to conceive the project. We thank Judith Lutton who has provided valuable insight into some of the computational challenges throughout the project. From the McAinsh lab we thank Muriel Erent for tissue culture training and initial creation of cell lines and Nina Pučeková for sample preparation. We thank Victoria Mitchell for her help in labelling training data for the model. We thank Hradini Konthalapalli from the Pines lab for their help in testing CelFDrive at the Institute of Cancer Research.

## 8 Funding information

This work was supported by the Medical Research Council (MRC) by iCASE Studentship [MR/R015910/1] project reference 2430216, T.B. was supported by funding from EPSRC (Engineering and Physical Sciences Research Council) [EP/V062522/1]. A.M., S.T-P. and D.S.C. are supported by a Wellcome Discovery Award [226605/Z/22/Z]. The Lattice light sheet microscope was established with a Wellcome Multi-user equipment grant[208384/Z/17/Z] to A.M.

## 9 Competing Interests

B. B. and K.K. are part-owners of 3i. The other authors declare no competing financial interests.

## Notes

https://github.com/scott-vision/CelFDrive

